# Reproductive and Seasonal Impacts on Rock Iguana Health: A Cross-sectional Study of Microbiome and Physiological Changes

**DOI:** 10.1101/2025.11.19.689195

**Authors:** Siefeldin E. Sobih, Erin L. Lewis, Karen Marie Kapheim, Susannah S. French

## Abstract

In this study, we examined how natural seasonal cycles and sex-based differences interact together using wild northern rock iguana as a study system. We collected samples during two seasons: reproductive (June) and post-reproductive (September) in 2016, this chapter demonstrates that sex and season both shaped microbiome composition. Reproductive females exhibited reduced microbial diversity during breeding compared to males and post-reproductive females, likely reflecting reproductive demands and dietary shifts. Physiological metrics including triglycerides, glucose, oxidative stress, and body mass index also influenced microbiome diversity, though the microbial community remained relatively stable across seasons.

## Introduction

The gut microbiome is the community of microorganisms inhabiting the digestive tract and has emerged as a fundamental component of animal biology (Hernández et al., 2024; Stothart et al., 2023). In some cases, these host-associated microbiomes allow their hosts to utilize nutrients that would be inaccessible without them (Q. Li et al., 2023), participate in immune function (Ki et al., 2024), and even affect behavioral and ecological fitness of their hosts (Gould et al., 2018). Studies have demonstrated that the gut microbiome is influenced by multiple factors, including diet, sex, age, and seasonal changes (Baniel et al., 2021; Kartzinel et al., 2019; Youngblut et al., 2019). However, our understanding of how seasonal fluctuations affect wildlife gut microbiome composition remains limited (Giacometti et al., 2024), particularly in the context of species conservation and wildlife health.

Seasonal changes are a recurring environmental factor that can significantly impact wildlife gut microbiomes (Stothart et al., 2023). Understanding the strength of the influence of seasonal variation and how it interacts with other factors remains challenging. This is largely due to a lack of comprehensive studies that examine both intrinsic and extrinsic influences in wild populations (Maurice et al., 2015). Seasonal fluctuations in the gut microbiome appear to be strongly influenced by diet. In wildwood mice, for example, seasonal differences in microbiome composition corresponded with a dietary shift from insects to seeds, and similar links between seasonal diet and microbiome changes have been observed in reptiles and wild gibbons (Maurice et al., 2015; Q. Li et al., 2023).

Similarly, intrinsic factors, such as host sex, can have a significant impact on the microbial composition. For instance, a study on the lizard species *Sceloporus virgatus* revealed significant sex-based differences in gut microbiota, with females exhibiting much lower diversity and richness compared to males (Martin et al., 2010). While sex remains a constant factor for many animals, its interaction with seasonal environmental changes or internal factors may influence microbiome dynamics in ways that are not yet fully understood (Comizzoli et al., 2021). Testing this hypothesis requires investigation of wildlife species experiencing seasonal variation, diet changes, and exhibiting sex-based differences in behavior, which could provide further insights into how those factors interact and influence gut microbiome composition.

Rock iguanas (*Cyclura cychlura*) in the Bahamas provide an excellent system for investigating the ecological and physiological influences of varying environmental factors. These large, primarily herbivorous lizards consume fruits, leaves, and flowers, and their diet shifts seasonally with available vegetation (Knapp et al., 2013). In addition, Northern Bahamian Rock Iguanas possess long lifespans, large body size, and robust metabolic, endocrine, and immune systems (French, Webb, et al., 2022a; Knapp et al., 2013). They occupy small, low-vegetation cays where they play a key ecological role in seed dispersal, nutrient cycling, and shaping vegetation (Knapp et al., 2013). Reproductive cycles are energetically demanding, with females producing 1-10 eggs annually and exhibiting marked hormonal shifts, while males show seasonal variation in testosterone (French, Webb, et al., 2022). This suggests that males and females may be influenced differently by seasonal changes, and these differences may correlate with changes to the microbiome and physiological aspects of health.

Additionally, seasonal and sex effects on the microbiome are potentially influenced by anthropogenic factors. In the Northern Bahamas, populations differ in exposure to tourism, with some receiving high-sugar or protein-rich foods from visitors, and others remaining largely undisturbed. Such anthropogenic supplementation has been linked to altered metabolism, immune responses, oxidative stress, and parasite loads, producing distinct physiological profiles (French, Webb, et al., 2022; Knapp et al., 2013). The combination of seasonal dietary changes, reproductive demands, sex-specific behaviors, and variability in human disturbance makes Rock Iguanas a powerful system for studying how different natural and anthropogenic factors shape the physiology and gut microbiome composition.

Here we explored seasonal variation in male and female rock iguanas to test three specific hypotheses. First, we hypothesized that the interaction between seasonal cycles and sex-specific behaviors shapes gut microbiome diversity, with microbiome composition fluctuating significantly between seasons and exhibiting unique patterns between males and females. Second, we hypothesized that rock iguana gut microbiome composition is influenced by host physiological factors, including hormonal changes and energy metabolism during reproductive cycles. We focused on two critically endangered subspecies, *C. c. inornata* and *C. c. figginsi*, across nine adjacent Exuma islands with varying levels of tourism exposure (high, moderate, and unvisited). In *C. c. inornata*, mating begins in early-mid May with nesting by mid-late June and females defending nests for 3-4 weeks after oviposition, while *C. c. figginsi* initiates breeding 1-2 weeks earlier. By September the post-reproductive phase marks the end of the mating season (French, Hudson, et al., 2022). Seasonal shifts in diet, hormones, energy metabolites, possible anthropogenic impacts, and behavioral changes during the mating season likely influence microbiome composition, and this study examines those interactions to add to our knowledge on how best to conserve these endangered iguanas.

## Methods

We analyzed 233 rock iguana samples that were sampled in 2016 from six islands in the Exumas during two seasons. Islands were classified into high, moderate, or low tourism based on the daily average number of tourists, and seasons were categorized as breeding in June and post-breeding for September (Table 3).

**Table 3.** Distribution of Tourism, Sex, and Season Across the Subspecies

| Tourism | Sex | Season | *C. figginsi* | *C. inornata* |
| --- | --- | --- | --- | --- |
| High | Female | Breeding | 7 | 15 |
|  |  | Post-Breeding | 14 | 14 |
|  | Male | Breeding | 12 | 12 |
|  |  | Post-Breeding | 12 | 15 |
| Moderate | Female | Breeding | 0 | 6 |
|  |  | Post-Breeding | 0 | 16 |
|  | Male | Breeding | 0 | 13 |
|  |  | Post-Breeding | 0 | 13 |
| Low | Female | Breeding | 18 | 3 |
|  |  | Post-Breeding | 9 | 10 |
|  | Male | Breeding | 11 | 4 |
|  |  | Post-Breeding | 16 | 13 |

### Study Design

Adult iguanas (snout-vent length >24 cm) were randomly sampled using hand capture, dip nets, or snares attached to a retractable pole (French, Webb, et al., 2022). For each subspecies, a trio of ecologically similar cays was selected that differ in human visitation levels: one heavily visited, one moderately visited, and one not visited. These sites facilitate direct comparisons of disturbance effects, with visitation intensity quantified by the number of boats and tourists. Genetic evidence indicates limited gene flow among populations (French, Webb, et al., 2022). On high- and medium-visitation islands, iguanas are frequently fed a variety of foods including lettuce, grapes, and other leftovers by tourists. Previous research has shown that these iguanas tend to be larger and heavier, but also display signs of dietary stress, including loose feces, elevated parasite loads, and altered blood chemistry (French, Hudson, et al., 2022).

In 2016, both species were sampled across high, moderate, and low tourism sites, but their distributions differed. *C. figginsi* was absent under moderate tourism and sampled only under high and low tourism, whereas *C. inornata* was sampled in all three conditions, with higher sample numbers overall (Table 3)

### Physiological and Immunity metrics

Blood samples were drawn from the caudal vein using a 1-1.5 ml syringe to assess key health indicators following the procedures described in Webb et al (2019). Similarly, plasma energy metabolites like triglycerides and free glycerol, reactive oxygen metabolites and antioxidant capacity were also assessed using the procedures described in Webb et al (2019). Innate immune function was assessed using a bacterial killing assay, which evaluates the capacity of blood plasma to eliminate *Escherichia coli* through the combined actions of immune proteins, antimicrobial peptides, and natural antibodies, as described in French, Webb, et al (2022). Energy metabolites, oxidative stress, and bacterial killing ability protocols followed those as described in French, Webb, et al (2022), with the modification that absorbance wavelength was read on a Byonoy spectrophotometer (Absorbance 96; Byonoy; Hamburg, Germany). Body mass index was calculated by dividing the lizard’s mass by the square of its snout-vent length (McCaffrey et al., 2023). Corticosterone, an indicator of stress, was measured along with other hormones, including progesterone and estrogen in females, and testosterone in both sexes (French, Hudson, et al., 2022; Ki et al., 2024). A total of 513 physiology samples were available before any data cleanup.

### DNA extraction

To sample gut microbial communities composition, iguanas from both subspecies were captured across islands with varying levels of tourism. Samples were collected by gently inserting sterile swabs through the vent into the cloaca and rotated for several seconds. After removal, the swab tip was broken off into a sterile 1.5 ml microcentrifuge tube, placed immediately on ice, and later stored at -20°C. Once swabs were sent back to the lab they were stored at -80°C until extraction (Kim et al., 2017). Genomic DNA was extracted from cloacal swabs using DNeasy PowerSoil Kits (QIAGEN, 12888-100) according to the manufacturer’s protocol. DNA yield was measured with a Qubit 2.0 Fluorometer using the High Sensitivity assay. Extracted DNA was stored at -80°C until sequencing. Negative controls, including extraction blanks and PCR no-template controls (NTCs), were processed alongside samples to monitor contamination (Ki et al., 2024).

### Sequencing

Bacterial DNA amplification and sequencing followed the protocol described by Ki et al (2024). For amplification, the V4 region of the 16S rRNA gene was amplified using primers 515F (Parada et al., 2016) and 806R (Walters et al., 2016) with Illumina-specific adapters and 12 bp Golay barcodes. PCR products were verified by gel electrophoresis, cleaned and normalized using SequalPrep Normalization Plate Kit, and pooled for paired-end sequencing on the Illumina MiSeq platform with PhiX Control library. We sequenced a total of 257 samples.

### Pre-processing and Feature Table

Demultiplexed FASTQ files from the 2016 sequence batch were imported into QIIME 2 (Amplicon 2025.4) (Bolyen et al., 2019). Primer sequences were removed using the cutadapt trim-paired command, with 515F (GTGYCAGCMGCCGCGGTAA) and 806R (GGACTACNVGGGTWTCTAAT) trimmed from the forward and reverse reads, respectively. Sequence quality was assessed using demux summarize, and low-quality bases were truncated using DADA2 (Callahan et al., 2016). Reads were truncated at 233 bp (forward) and 195 bp (reverse), corresponding to the position where the median quality score dropped below 30, with no trimming from the 5′ ends.

### SILVA Reference Database Processing and Taxonomic Classification

The RESCRIPt plugin (Robeson et al., 2021) in QIIME 2 was used to process the SILVA 138.2 NR99 database. First, the SILVA database was retrieved with species labels excluded and taxonomic rank propagation enabled. RNA sequences were reverse-transcribed to DNA format and quality-filtered. Sequences were then length-filtered by domain, with minimum lengths set to 900 bp for Archaea, 1,200 bp for Bacteria, and 1,400 bp for Eukaryota. The filtered sequences and their corresponding taxonomy were dereplicated using the lowest common ancestor (LCA) method. To match the amplified region, primer-specific reads were extracted, targeting the 515F and 806R primer regions and retaining fragments between 100 and 400 bp. These reads were dereplicated again using RESCRIPt and used to train a Naive Bayes classifier. Taxonomic classification was performed on representative sequences.

### Phylogenetic Tree Construction and Data Export

Multiple sequence alignment of representative sequences was performed using MAFFT and subsequently masked to remove hypervariable regions. A phylogenetic tree was then constructed using FastTree through the phylogeny plugin in QIIME 2, generating both rooted and unrooted trees for downstream phylogenetic analyses. Taxonomy, rooted tree, and feature table output files were exported. The final output contained 255 samples and 8,362 taxa.

### Contaminant and Low-Abundance Feature Filtering

The initial quality-filtered feature table, rooted tree, and metadata were imported into an *R (version 4.5.1)* environment as a phyloseq object using the *phyloseq (version 3.21)* package for further downstream quality control and analysis (McMurdie & Holmes, 2013). A fixed random seed was set to ensure reproducibility. Next, taxa classified as mitochondria (Family level) and chloroplasts (Order level) were removed. To identify potential contaminants, negative control samples were flagged, using the *decontam (version 3.21)* package (Davis et al., 2018) with the prevalence method set to a threshold of 0.5. A total of 70 contaminant features were identified and removed. Control samples were then excluded from further analyses. Taxa lacking phylum-level classification were removed.

A prevalence filter was applied as described in Nearing et al (2022a). Within each sample, ASVs with counts fewer than 10 were set to zero. ASVs that had a total count of zero across all samples after this filtering were excluded from the dataset. This retained 3,960 taxa. Finally, samples with fewer than 1,000 total reads were excluded, leading to a final dataset of 237 samples. The filtered phyloseq was used for further downstream analyses.

### Multicollinearity and Model Parameters

Prior to analysis, we tested for multicollinearity among continuous predictor variables using the Variance Inflation Factor function from the *car (version 3.1-3)* package. Variables with perfect collinearity (GVIF = 100%) were removed. For the remaining predictors, GVIF^(1/(2Df)) values were evaluated, and variables with values exceeding ∼2.5 were considered highly collinear. When multiple variables exhibited high collinearity (e.g., mass, snout-vent length (SVL), and body mass index), the most biologically relevant or data-complete variable was retained (e.g., body mass index was included as a representative covariate in the analysis; mass and SVL were not included). Because some sex steroids were only available in female iguanas (progesterone and estrogen), we ran separate statistical analyses for them. We omitted variables with missing data (NA) greater than 15% across all samples.

The final set of continuous covariates included: glucose, testosterone, corticosterone, bacterial killing ability, glycerol, triglycerides, dROMS, antioxidant index, and body mass index. Categorical factors included: tourism category, sex, and reproductive season across a total of 237 samples. The unique id column was used to account for repeated measures as a random effect. Additionally, progesterone and estrogen were retained in the female-specific models. Finally, continuous covariates were standardized using z-score transformation and all the graphs were created using the ggplot2 package in *R*.

### Relative Abundance

To visualize microbial community composition for both the whole dataset and data ordered by different seasons and both sexes we used *microViz* package (D. Barnett et al., 2021). First, the phyloseq object was checked for errors in the taxonomic labels. Ambiguous classifications were reassigned to a higher taxonomic rank, and unknown taxa were assigned to *Incertae Sedis*. Next, microbial communities were aggregated at the phylum level, then ordered based on Bray-Curtis distance and Ward’s clustering method applied to compositionally transformed data. Then we ran a separate abundance analysis faceted by season * sex, and aggregated on the family-level taxonomy. Finally, taxa were filtered to have a minimum abundance of 2% reads in one or more samples. Bar plots were generated with colors for top 10 phyla or family, with less abundant ones grouped under “others”.

### Alpha Diversity Statistical Analysis

Faith’s phylogenetic diversity (PD) was calculated using the *calculatePD()* function on the rooted phylogenetic tree, and Shannon was computed using *estimate_richness()*. Linear mixed models assessed the effects of continuous and categorical covariates on alpha diversity metrics with unique id as a random effect for repeated measures. PD and Shannon were Box-Cox transformed, with the optimal λ selected automatically, to satisfy model assumptions (homoscedasticity and normality of residuals) and minimize AIC among other transformation options tested. The mixed models were passed to the *step()* backward elimination function from the *lmerTest* package to choose the best model, and in some cases it eliminated the random effect and recommended a simple linear model.

Significance testing was performed via Type II ANOVA from the *car* package in *R*, with White adjustment. Pairwise comparisons of estimated marginal means were evaluated using the *emmeans* package. For each diversity metric, null models (intercept-only) were compared to models with covariates, to confirm the contribution of explanatory variables.

### Community Composition Ordination

Beta diversity was assessed using Bray-Curtis dissimilarity distance. Ordinations of Principal Coordinates Analysis (PCoA) were calculated to visualize the compositional distribution in 2D space, using the *phyloseq* package. Bray-Curtis distances were transformed to log-transformed data using a log(1 + x), which retains zero values while stabilizing variance. Ordination results were visualized separately for each sub-species, sex within reproductive season, and tourism categories.

### Community composition and beta diversity

Microbial community composition differences were assessed using PERMANOVA (Permutational Multivariate Analysis of Variance) using the *adonis2* function in the *vegan (version 2.7-0)* package (Oksanen et al., 2025). Analysis was conducted on the log(1+x) transformed Bray-Curtis distance for each sub-species. The by = "margin", by = "terms", and by = NULL options were used to evaluate marginal, term-wise, and full model effects, respectively. A total of 9999 permutations were used in all tests and *P*-values adjusted for multiple comparisons with the Benjamini–Hochberg (BH) method. Next, post hoc pairwise comparisons were performed using the *pairwise.adonis* function. The comparisons were conducted for tourism level and reproductive score, with *P*-values adjusted for multiple comparisons using the Benjamini-Hochberg (BH) method.

Because PERMANOVA assumes homogeneity of variance within variables, differences in dispersion were tested using the *betadisper* function. This was done for tourism categories, reproductive season, sex, and subspecies. Significance was assessed using *anova()* and pairwise group comparisons were evaluated using *TukeyHSD()* function.

### Differential Abundance Analysis

Differential abundance analysis with bias correction was done using ANCOM-BC2 (Lin & Peddada, 2024) at the “Family” taxonomic level on the phyloseq object. A contrast variable of sex * reproductive season was used as part of the fixed-effect formula, and selected for pair-wise comparison. For each analysis run, detecting structural zeroes was enabled for the detection of taxa that are completely absent in one group but present in another (Lin & Peddada, 2024). Pseudo-sensitivity analysis was also enabled to assess robustness of differential abundance to the presence of pseudo-counts (Lin & Peddada, 2024). Consistent with our earlier filtration steps, only samples with 1000 reads were retained for this analysis, and taxa were included only if they are present in 10% of the samples. A small constant of 0.05 was added to stabilize variance estimates. Also, the direct comparisons were run against females in the breeding season as a reference group. Finally, *P* values were adjusted using the built-in Benjamini-Hochberg (BH) method with a significance threshold of *α* = 0.05.

## Results

A total of 233 samples of two rock Iguana sub-species were analysed. After filtering, 6,820,366 total high-quality reads were retained, with an average of 29,272 ± 625 reads per sample. After error correction and dereplication, high-quality reads were denoised into 3,960 ASVs. Unique taxonomic identification of ASVs resulted in a total of 45 phyla, 81 classes, 166 orders, 270 families, 585 genera, and 585 bacterial species.

### Fecal Microbiome Composition of Rock Iguana across Reproductive Season and Sex

#### The Relative Abundance of Microbial Composition

The 10 most dominant Phyla were Bacillota (36.64%), Actinomycetota (32.44%), Campylobacterota (10.10%), Bacteroidota (7.97%), Pseudomonadota (4.32%), Verrucomicrobiota (1.52%), Methanobacteriota (1.47%), Spirochaetota (1.12%), Cyanobacteriota (0.97%), and Planctomycetota (0.78%), representing 97.33% of the total sequences (Figure 9).

**Figure 9:**
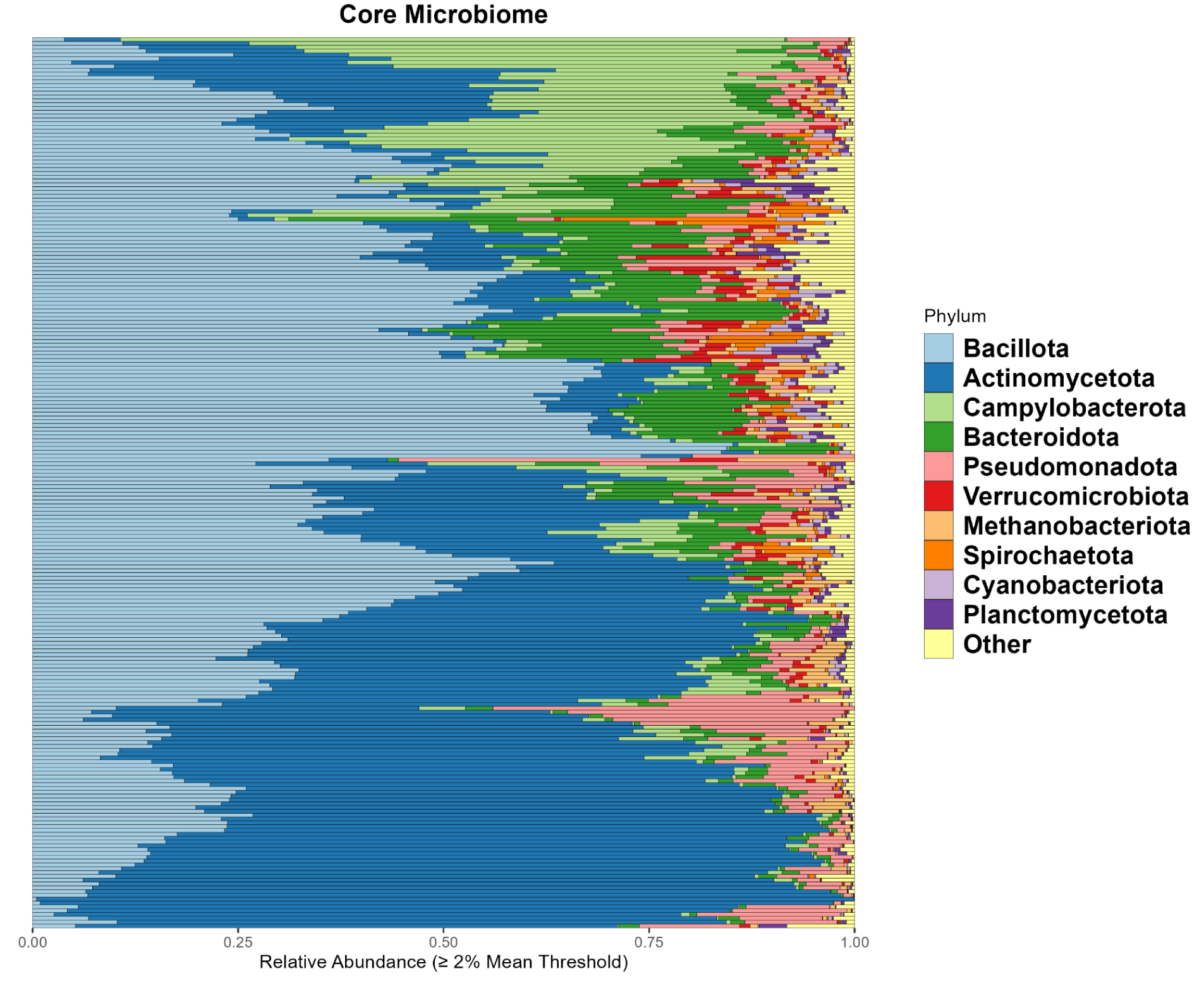
Hindgut Microbiome Phylum-Level Composition in Rock Iguanas Hindgut microbiome composition of rock iguanas. Phylum-level relative abundance present at ≥ 2% in at least one sample. Each horizontal bar represents one sample (n = 233), with relative abundance values ranging from 0.00 to 1.00 on the x-axis. The top 10 most abundant phyla are shown in distinct colors: Bacillota (light blue), Actinomycetota (dark blue), Campylobacterota (light green), Bacteroidota (medium green), Pseudomonadota (light red), Verrucomicrobiota (red), Methanobacteriota (light orange), Spirochaetota (orange), Cyanobacteriota (purple), and Planctomycetota (dark purple). Less abundant phyla are grouped together and shown in yellow as "Other."

To investigate our primary hypothesis that the Rock Iguana gut microbiome diversity fluctuates across different seasons, with unique patterns between males and females, we examined the relative abundance profiles of bacterial families during the breeding and post-breeding seasons, stratified by sex. These profiles showed broadly similar community patterns (Figure 10). In both seasons, Corynebacteriaceae, Helicobacteraceae, and Lachnospiraceae were dominant, together accounting for over 30% of the total family-level abundance.

**Figure 10:**
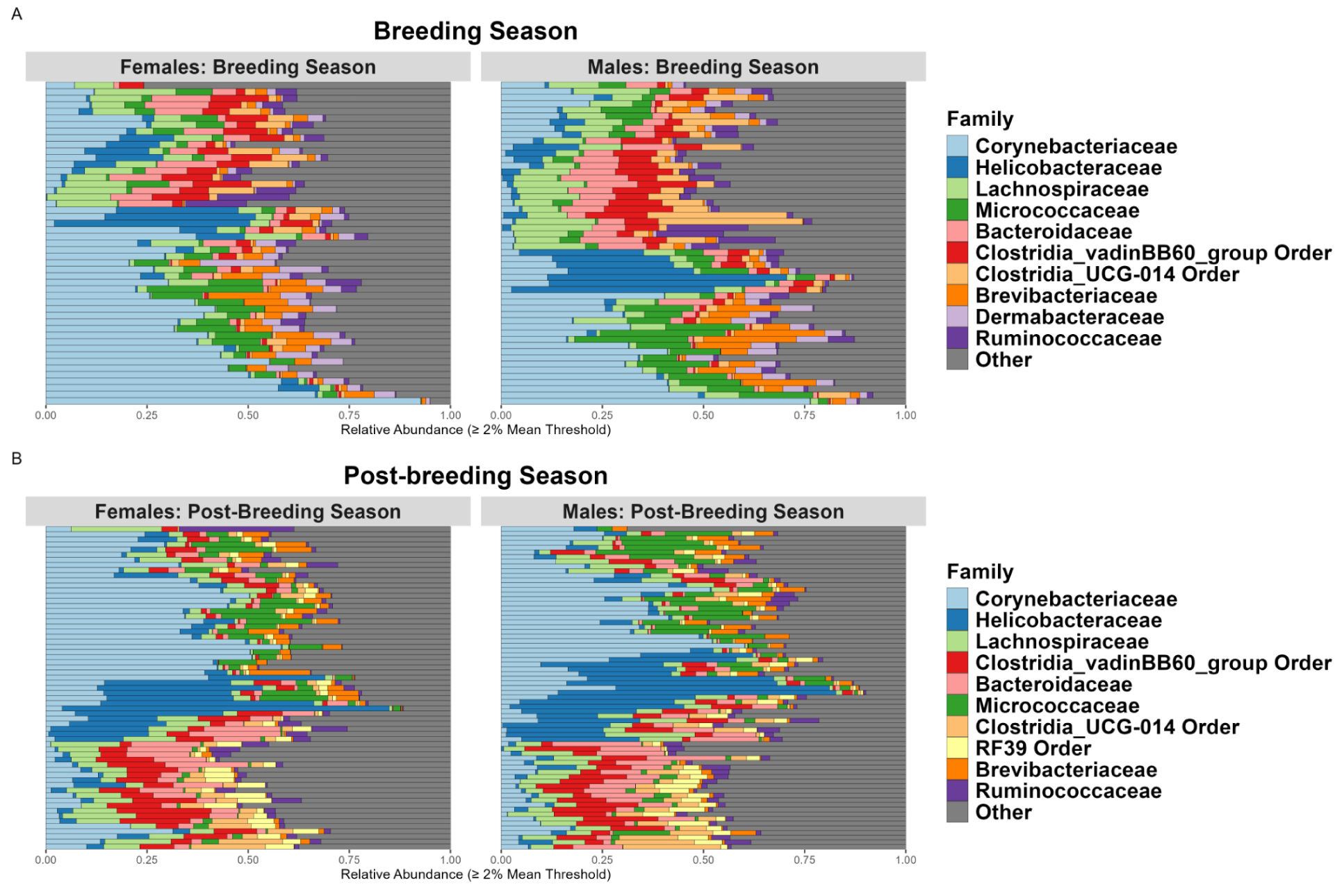
Relative Abundance of Dominant Bacterial Families in *C. Inornata* and *C. Figginsi* Across Reproductive Seasons and Sex Relative abundance of bacterial families of breeding (A) and post-breeding (B) grouped by sex. Each horizontal bar represents a single sample. Only the top 10 most abundant families were colored, the rest were grouped and colored under “Other”.

#### The Differential Abundance of Microbial Composition

We identified ten microbial families that were significantly differentially abundant between sex and reproductive season categories, but only one passed the sensitivity threshold (Figure 11). Specifically, five taxa showed significantly reduced relative abundance in breeding males compared to breeding females (Figure 11A): Staphylococcaceae (*lfc = -2.69, P < 0.05*), Steroidobacteraceae (*lfc = -2.71, P < 0.05*), Nitrososphaeraceae (*lfc = -2.87, P < 0.05*), Eubacteriaceae (*lf = -3.09, P < 0.05*), and Dermacoccaceae (*lfc = -7.71, P < 0.05*). Post-breeding males had significantly lower abundance of three taxa compared to breeding females: Alteromonadaceae (*lfc = -2.95, P < 0.05*), Geminicoccaceae (*lfc = -3.01, P < 0.05*), and Dermacoccaceae (*lfc = -10.40, P < 0.05*). However, Coxiellaceae (*lfc = 5.71, P < 0.05*) was significantly higher in post-breeding males compared to breeding females (Figure 11C). Notably, Dermacoccaceae was significantly less abundant in all other groups when compared to breeding females (breeding males: *lfc = -7.71, P < 0.05*; post-breeding males: *lfc = -10.40, P < 0.05*; post-breeding females: *lfc = -8.49, P < 0.05*; Figure 11).

**Figure 11:**
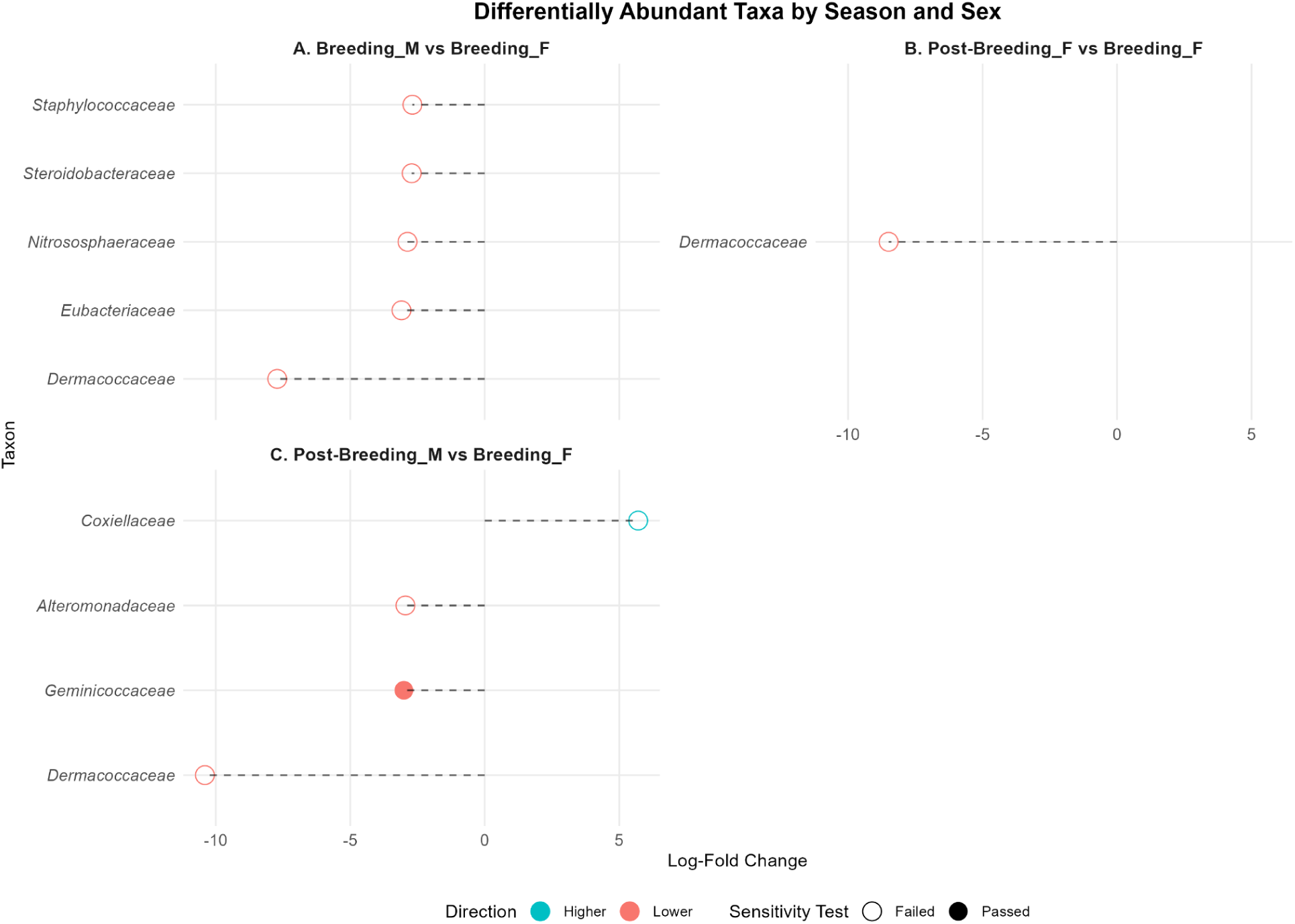
Differential Abundance of Bacterial Families on Breeding and Post-breeding Seasons in Both Male and Female Iguanas Each panel compares a combination of males/females and post-breeding/breeding abundances. Taxon names are on the y-axis, and log-fold change is on the x-axis. Horizontal dotted lines extend from zero to each point, representing a taxon’s log-fold change. Point color indicates whether a taxon increased (blue) or decreased (red) in comparison to the breeding season females (reference group), and the dot color fill indicates whether it passed (filled) or failed (empty) the sensitivity analysis.

### Shifts in Community Composition and Alpha Diversity Across Reproductive Season and Sex in Rock Iguanas

#### Changes in Alpha Diversity

Faith Phylogenetic Diversity (PD) and Shannon were used to determine which factors affected alpha diversity patterns. For PD, the best-fitting model was a significant improvement over the null model (*F = 6.01, P < 0.0001*) and included species, tourism level, BMI, sex, reproductive season, and the interaction between sex and reproductive season. PD was best predicted by the interaction between sex and reproductive season (*F = 6.09, P = 0.014*; Figure 12A), indicating that the influence of sex on diversity varied across reproductive periods. Subspecies (*F = 8.70, P = 0.036*; Figure 12C) and tourism (*F = 3.59, P = 0.029*; Figure 12B), and higher Body Mass Index (BMI) (*F = 4.74, P = 0.031*; Figure 12B) was likewise associated with altered alpha diversity patterns. Tukey-adjusted pairwise comparisons revealed that males in the breeding season had higher PD than females in the breeding season (*t =* -2.769, *P* < 0.05; Figure 12A). All other combinations of sex and reproductive season interactions were not significant (*P > 0.05*). Among tourism categories, PD was significantly higher on high-tourism islands compared to no-tourism islands (*t = 2.57, P = 0.030*; Figure 12B), while other pairwise comparisons were not significant (*P > 0.05*). As for subspecies, *C. Inornata* had higher PD in comparison to *C. figginsi* (*t =* -3.430, *P* < 0.01; Figure 12C).

**Figure 12:**
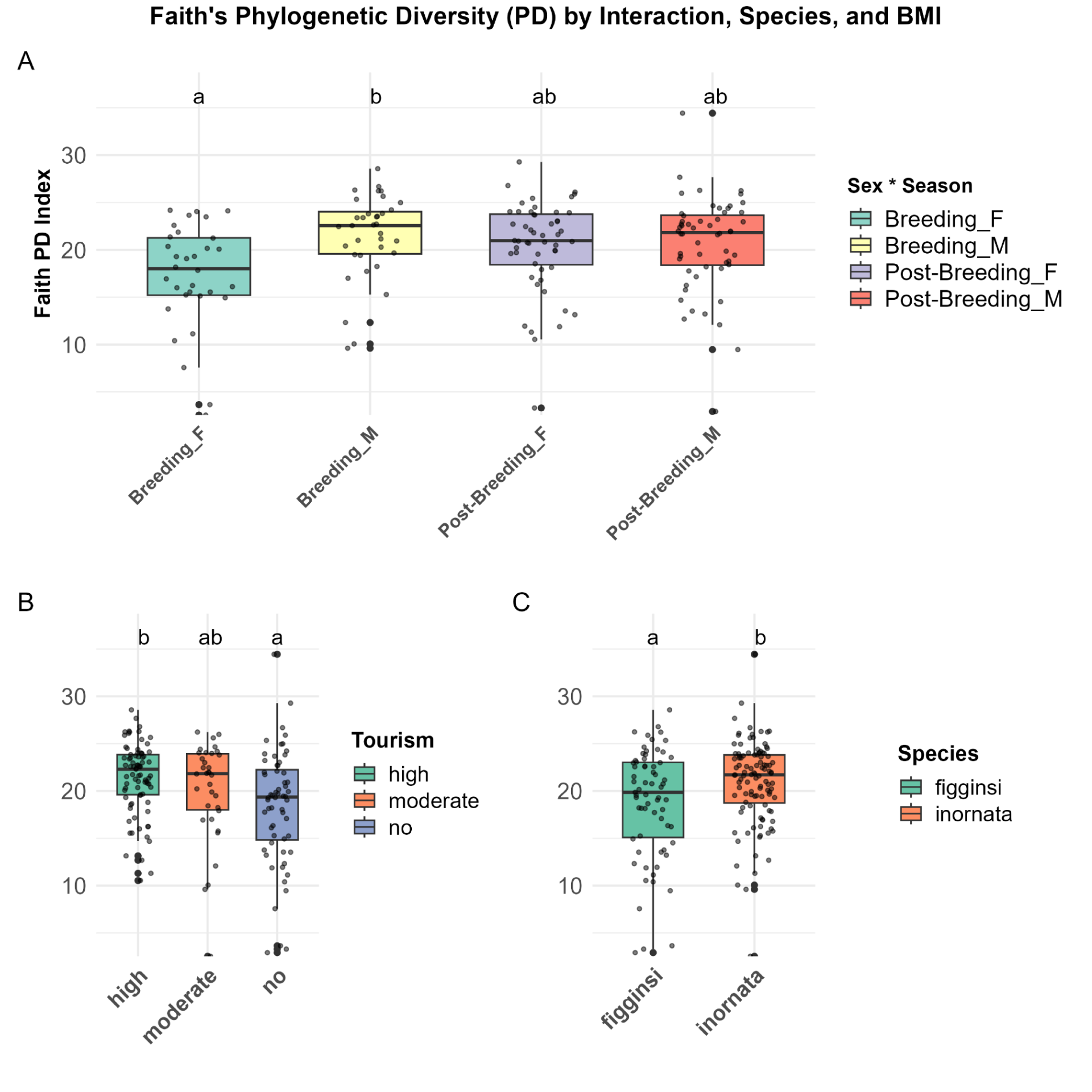
Faith’s Phylogenetic Diversity (PD) Across Reproductive Season * Sex, Tourism, and Species Faith’s Phylogenetic Diversity (PD), (A) PD across reproductive season * sex. (B) PD across tourism categories. (C) PD across species. Different letters indicate significant pairwise differences (Tukey-adjusted, *P* < 0.01). Boxes show interquartile range; horizontal lines denote medians; dots represent individual data points.

PD was assessed in females only with progesterone and estrogen included in the model to evaluate their influence on alpha diversity. In this model, PD was best predicted by species (*F = 11.43, P = 0.001*), glucose levels (*F = 7.25, P = 0.0088*), and reproductive season (*F = 14.87, P = 0.00025*). Also, dROMS showed a significant effect (*F = 14.04, P = 0.00036)*, However, neither progesterone nor estrogen had any significant impact on PD.

As for the Shannon index, the best-fitting model improved on the null model (*F = 4.7, P < 0.001*) and included species, triglycerides, sex, reproductive season, and the interaction between sex and reproductive season. Shannon index was best predicted by the interaction between sex and reproductive season (*F = 6.50, P = 0.012;* Figure 13A), species (*F = 5.76, P = 0.018;* Figure 13B) and triglycerides (*F = 5.17, P = 0.024*). Tukey-adjusted pairwise comparisons showed that males in the breeding season had higher Shannon diversity than females in the breeding season (*t = -2.563, P < 0.05*; Figure 13A). In addition, both females (*t = -2.076, P = 0.02*; Figure 13A) and males (*t = -2.184, P = 0.011*; Figure 13A) in the post-breeding season had higher Shannon diversity than females in the breeding season. All other combinations of sex and reproductive season interactions were not significant (*P > 0.05*). Subspecies, *C. Inornata* had higher Shannon diversity in comparison to *C. figginsi* (*t =* -1.14, *P* = 0.018; Figure 13B).

**Figure 13:**
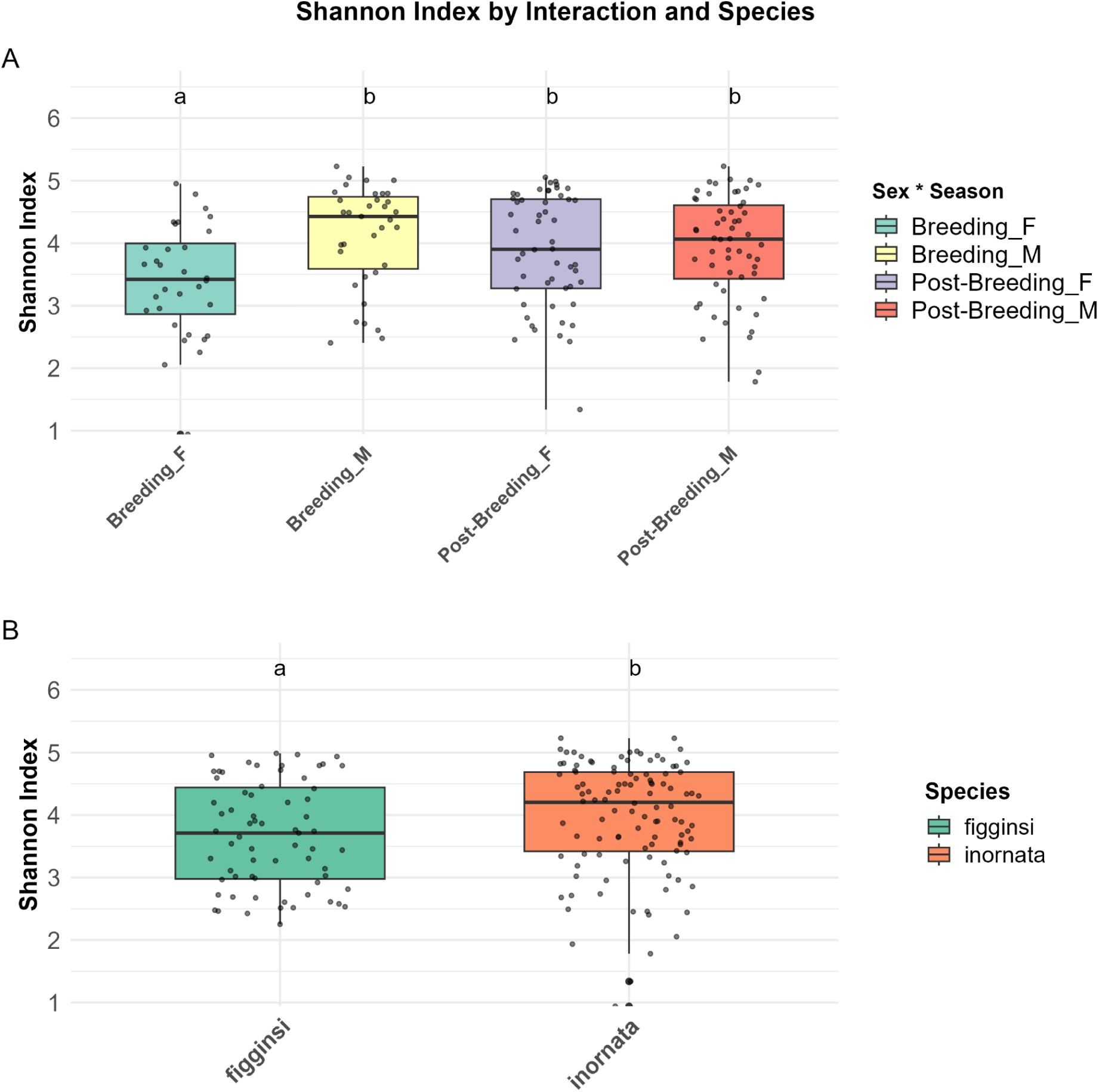
Shannon Across Reproductive Season * Sex and Species (A) Shannon index across reproductive season * sex. (B) Shannon index across species. Different letters indicate significant pairwise differences (Tukey-adjusted, *P* < 0.01). Boxes show interquartile range; horizontal lines denote medians; dots represent individual data points.

Like PD, the Shannon index was modeled in females with progesterone and estrogen included to evaluate their influence on alpha diversity. We found that species (*F = 6.44, P = 0.013*), glucose (*F = 9.62, P < 0.01*), reproductive season (*F = 17.32, P < 0.001*), and dROMS (*F = 13.00, P < 0.001)* were all significant predictors of Shannon variability. Neither progesterone nor estrogen had a significant impact on Shannon.

#### Microbiome Community Composition and Ordination

To assess variation in microbiome composition between reproductive seasons and sex, we conducted PCoA on Bray-Curtis distances (log(1+x) transformed; Figure 14). In tourism and reproductive groups, no clear separation was observed (Figure 14A; 14B). However, subspecies showed a distinct clustering along axis 2 (Figure 14C).

**Figure 14:**
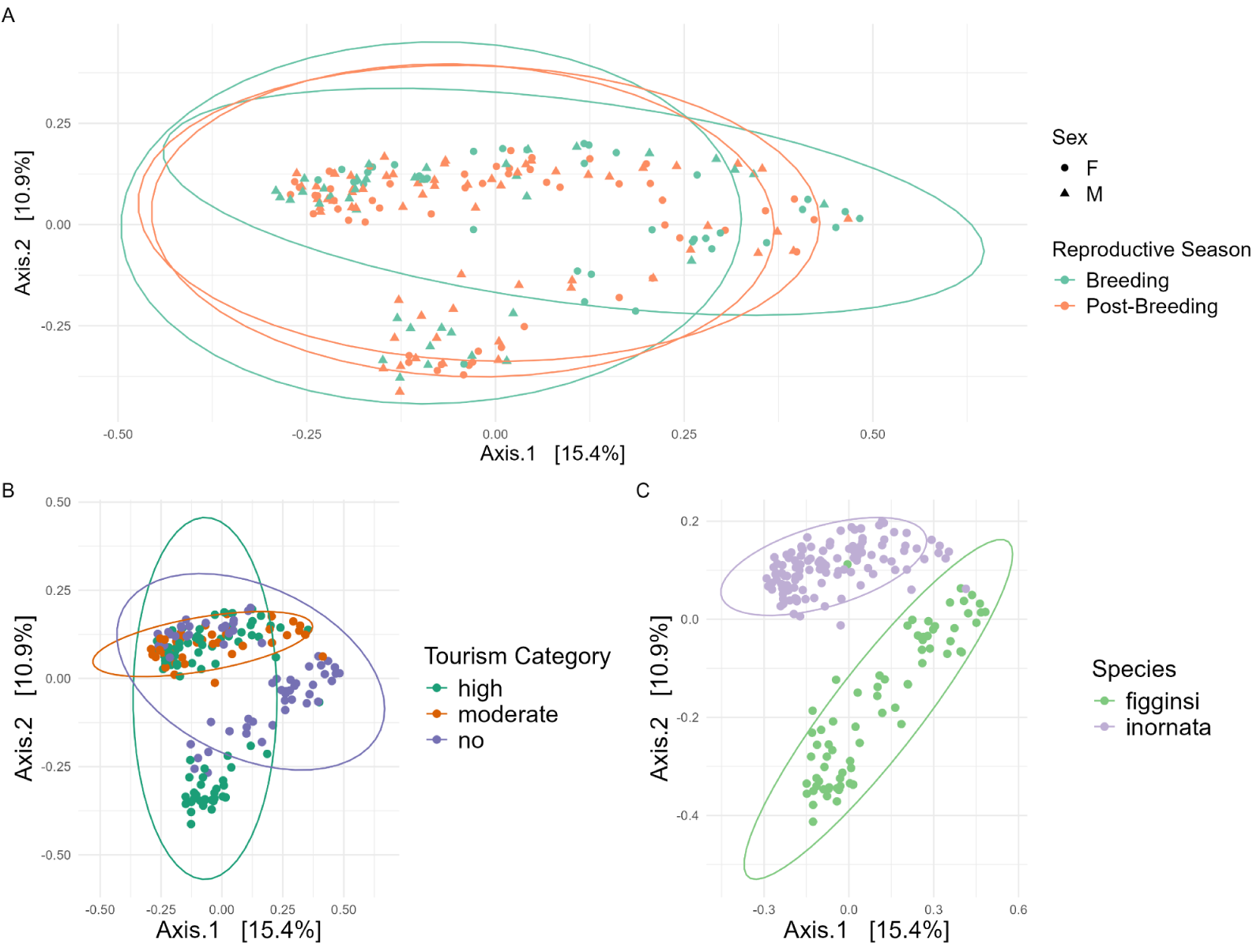
Principal Coordinates Analysis of Log(1+x)-Transformed Bray-Curtis Dissimilarities on Rock Iguana Microbial Communities. Principal Coordinates Analysis (PCoA) of log(1+x)-transform Bray-Curtis dissimilarity matrix. Each point represents an individual iguana sample. (A) Samples colored by season and shaped by sex (circles = females and triangles = males). (B) Samples colored by tourism category. (C) Samples colored by subspecies. Axis indicate the proportion of variance explained by each principal coordinate. Ellipses for each figure were done using multivariate t-distribution.

PERMANOVA analysis revealed that multiple factors significantly impact community structure. The model explained 25% of variance (R² = 0.25, *P* < 0.001), with significant marginal predictors including species (R² = 0.071, *P* < 0.001), tourism category (R² = 0.039, *P* < 0.001), glycerol (R² = 0.007, *P* = 0.049), triglycerides (R² = 0.007, *P* = 0.0299), body mass index (R² = 0.009, *P* = 0.0068), and the reproductive season * sex interaction (R² = 0.007, *P* = 0.0431). In addition to the marginal predictors, the analysis of term effects, which represents the unique contribution of each factor in the model, showed that sex (*R²* = 0.009, *P < 0.01*) and reproductive season (*R²* = 0.019, *P < 0.001*) significantly impacted community composition.

Pairwise PERMANOVA with Benjamini-Hochberg correction revealed that all tourism pairs (high vs moderate, high vs no, and no vs moderate) and species (*C. inornata* vs *C. figginsi)* are all significant (*P < 0.001)*. Similarly, all pairwise comparisons for season * sex interactions were significant (*P < 0.01*), except for post-breeding season males vs females (*P < 0.83)*.

To evaluate the impact of group dispersion on community composition, we tested for homogeneity of multivariate dispersion. Dispersion differed significantly in the tourism variable (*P < 0.001;* Figure 15A), with significant differences between the no tourism category vs moderate and high tourism islands (*P < 0.001;* Figure 15A). Similarly, there was a significant difference in group dispersion between males and females (*P = 0.045;* Figure 15B), and between both subspecies (*P < 0.001;* Figure 15D). However, only the reproductive season did not have significant differences in group dispersion (Figure 15C).

**Figure 15:**
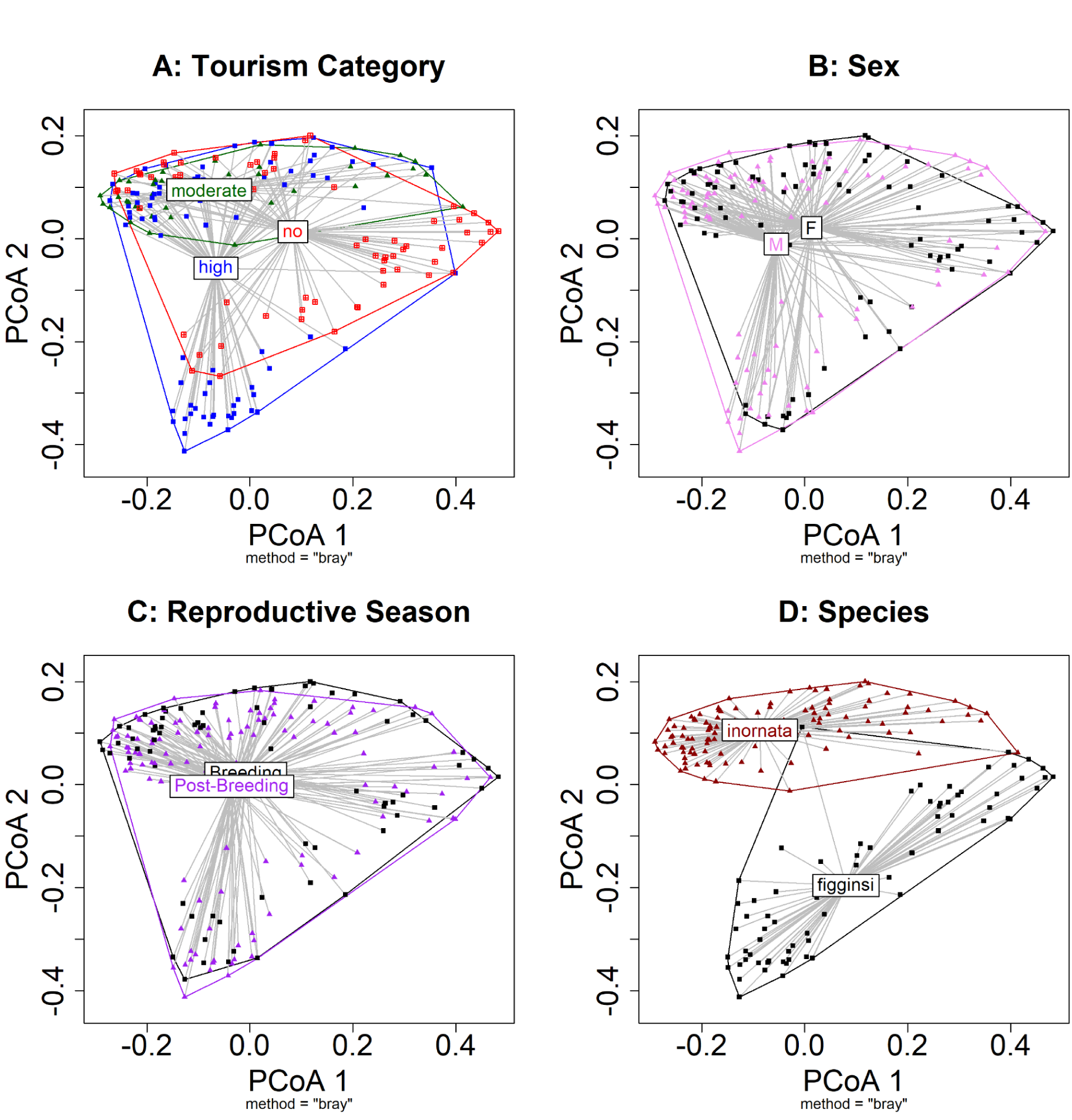
Beta Dispersion of Rock Iguana Samples Based on Log(1+x)-Transformed Bray-Curtis Dissimilarities Across Subspecies, Tourism, Sex and Season. Beta dispersion plots based on log(1+x) transformed Bray-Curtis dissimilarity metric. Distances to group centroids are shown for (A) Tourism, (B) Sex, (C) Reproductive Season, and (D) Species. Each point represents an individual sample, with lines connecting samples to their group centroid. Polygons encompass the spatial distribution of each group in ordination space.

## Discussion

In this study we examined the influence of reproductive seasonal variation and sex on the rock iguana gut microbiome, we compared samples collected across two reproductive seasons in 2016 from six populations of rock iguanas. This comparison allowed us to test our hypotheses: (1) that the rock iguana gut microbiome diversity fluctuates significantly between different seasons, with unique patterns between males and females, and (2) that the rock Iguana gut microbiome composition is influenced by the physiology of the host. Our results supported these hypotheses: season and sex both shaped microbial composition, females exhibited reduced diversity during reproduction, and host physiological metrics such as triglycerides, glucose, d-ROMS, and BMI were associated with microbial composition.

### Season, Sex, and Diet Drive Shifts in Rock Iguana Microbiome Composition

Seasonal and sex-specific reproductive demands appear to be major drivers of microbiome dynamics in rock iguanas. Specifically, the interaction between season and sex explained a significant portion of the variance in both alpha and beta-diversity, with certain microbial families increasing or decreasing depending on sex across seasons. Females showed notably reduced alpha diversity during the breeding season compared to breeding males, post-breeding females, and post-breeding males (Figure 12). This pattern likely reflects the physiological demands of gravidity and nesting, when energy allocation shifts and dietary options may change (French, Webb, et al., 2022). Similar patterns were seen in other *sceloporus virgatus*, where microbiome composition shifted with season and sex, and diversity was lowest in females during reproduction and periods of increased male-female social interactions (Bunker, Arnold, et al., 2022). These changes were attributed to several interacting factors: increased intersexual contact during breeding leading to the enrichment of already dominant taxa (such as Enterobacteriaceae), stress and hormonal fluctuations in steroid hormones associated with reproduction, and sex-specific behavioral differences in activity and food intake. In gravid females, microbiome remodeling may reflect selective pressures to protect eggs and shape offspring development. Comparable patterns occur in other iguanian lizards: female *Sceloporus virgatus* transfer a high quantity, highly selective microbiome to their eggshells, which suppresses fungal growth (Bunker et al., 2021). This vertical transmission appears to be a significant mechanism enhancing female reproductive fitness (Bunker et al., 2021).

Human activities, particularly tourism, significantly influence the gut microbiome of rock iguanas by introducing anthropogenic food sources that affect both alpha and beta diversity (French, Webb, et al., 2022; Knapp et al., 2013). This plasticity of gut microbiomes in response to dietary changes is demonstrated in a controlled study on non-reproductive omnivorous lizards, *Liolaemus ruibali*, which showed that diet manipulation alone induced significant changes in gut morphology and microbiome composition (Kohl et al., 2016). Together, these findings suggest that dietary shifts from human activities significantly influence gut microbiome composition, with similar plasticity demonstrated in other lizard species making diet a key driver of variance in alpha- and beta-diversity.

### Rock Iguana Gut Microbiome Taxonomy Is Relatively Stable Across Seasons and Similar to Other Reptiles

Across both seasons and sexes, the microbial community was dominated by a consistent set of phyla (Figure 10), including Bacillota, Actinomycetota, Campylobacterota, Bacteroidota, and Pseudomonadota, along with families such as Corynebacteriaceae, Helicobacteraceae, and Lachnospiraceae. A similar pattern was found in previous work on lizards (Bunker, Martin, et al., 2022; Gilbert et al., 2017a; Hoffbeck, Middleton, Nelson, et al., 2025; Hoffbeck, Middleton, Wallbank, et al., 2025). In *P. lilfordi* and *P. pityusensis*, Bacillota (also called Firmicutes) constituted an average of 45.1% of the faecal microbiota (Alemany et al., 2023). It was similarly identified as the second most dominant phyla in several Portuguese lizard species (Vasconcelos et al., 2023). This phylum is especially significant in omnivorous and herbivorous reptiles, as taxa within the Lachnospiraceae family contribute to the breakdown of complex plant material and play a central role in fibre digestion (Alemany et al., 2023). The presence of these cellulolytic taxa aligns with the role of Firmicutes in shaping reptilian diet, as also observed in *Sceloporus virgatus*, where they comprised 62% of faecal communities (Hoffbeck, Middleton, Nelson, et al., 2025). In another study on *Sceloporus virgatus*, Lachnospiraceae was amongst the most abundant taxa recovered from faecal pellets, making up 38.7% on average, contributing to plant matter degradation (Bunker, Martin, et al., 2022).

Pseudomonadota (formerly Proteobacteria) has also been reported as among the most abundant phyla in other reptile microbiome studies. For example, it was among the dominant phyla in five lizard species (Alemany et al., 2023) and in *Sceloporus virgatus*. Its prevalence in the cloaca, particularly members of the family Enterobacteriaceae, has been linked to a specialized role in transferring antifungal bacteria to eggshells during oviposition, and enhancing egg survival (Bunker, Martin, et al., 2022).

Our differential abundance analysis revealed multiple families were more relatively abundant in breeding females than breeding males, post-breeding males, and post-breeding females. For example, the Staphylococcaceae family was enriched in breeding females to breeding males (Figure 11). Members of this family, particularly Staphylococcus, have been associated with reptiles experiencing stress (Ebani, 2024), which suggests a link between reproductive stress and microbial shifts. Other families that were prevalent in breeding females included Steroidobacteraceae, Nitrososphaeraceae, Eubacteriaceae, and Dermacoccaceae. Steroidobacteraceae are known for their ability to fix nitrogen and maintain this activity under fluctuating oxygen conditions in deadwood, as well as for their potential to produce diverse secondary metabolites (Richy et al., 2024). Rock iguanas may have acquired this family through incidental consumption of dirt or interactions with fallen barks, since they are common in decomposing wood and their relative abundance was low in both seasons (<0.5%). Nitrososphaeraceae, belonging to the Thaumarchaeota, are ammonia oxidizers converting ammonia to nitrite in terrestrial environments like soils (Thomas et al., 2022). This process is vital for mobilizing reactive nitrogen species and is a core genomic feature contributing to their energy metabolism in lizards (Vasconcelos et al., 2023). Eubacteriaceae, was less functionally defined, but have been reported as part of the core gut microbiota in reptiles (Baldo et al., 2018) and appear to vary seasonally in abundance, indicating possible dietary or seasonal associations (X. Zhu et al., 2024).

In contrast, post-breeding males displayed differences in Coxiellaceae, Alteromonadaceae, and Geminicoccaceae. Coxiellaceae contains one of its well-studied species *Coxiella burnetii*, a zoonotic pathogen that is usually associated with a tick that a reptile host is carrying (Mendoza-Roldan et al., 2021). Alteromonadaceae and Geminicoccaceae are often associated with environmental processes, like degrading algae (Kwak et al., 2012), or production of indole-3-acetic acid (Jiang et al., 2022), but none of them were linked to a functional role in reptiles. Dermacoccaceae consistently differed across all comparisons, especially post breeding females to breeding females, suggesting this family may be particularly sensitive to host reproductive status. This family was isolated from the amphibian and human skin with no functional or metabolic role in lizards (Flechas et al., 2017; Williams & MacLea, 2019). Dermacoccaceae likely vary because they are not native to the iguana gut (<0.5% relative abundance). Their fluctuations are more consistent with environmentally acquired, skin-associated bacteria responding to physiological and behavioral changes across reproductive stages, rather than gut-specific processes. Taken together, these differential abundance patterns suggest that the intensity of the reproductive season and sex-specific stress shape the microbiome, while a core community remains stable across seasons and sexes.

### Energy, Stress Markers, and Body Mass Index Significantly affect Gut Microbial Beta- and Alpha-Diversity

This study tested whether different hormones and physiological metrics influenced alpha diversity and microbial community structure in rock iguanas. We found that different physiological metrics affected males and females differently. While energy metabolites and body mass impacted microbial diversity in both sexes, female reproductive and stress hormones, such as estrogen, progesterone, and corticosterone, surprisingly showed no significant effects.

In female iguanas, we found that glucose and d-ROMS were significant factors associated with increased PD and Shannon indexes. In contrast, neither female hormones (estrogen, progesterone) nor the stress hormone corticosterone, which has been linked to energy and stress levels (French, Webb, et al., 2022) were significant predictors of alpha or beta diversity. Similarly, no other physiological variables were associated with beta diversity. One proposed mechanism for why glucose and d-ROMS were significant factors is the direct and indirect influence of the gut microbiota on redox reactions, which leads to changes in the inflammatory response (Kunst et al., 2023). A very similar pattern was also observed when a high-sugar diet in *Iguana iguana*, triggered an alteration to the immune function (Ki et al., 2024). This suggests that both d-ROMS and glucose levels are part of a larger mechanism connected to the gut microbiome that influences the immune response and may work bidirectionally.

In male iguanas, different variables affected different aspects of alpha diversity. Body mass index (BMI) and tourism primarily influenced phylogenetic diversity, suggesting effects on the evolutionary diversity of microbial lineages present. In contrast, triglycerides were associated with Shannon diversity, indicating shifts in both the richness and evenness of taxa within individual iguanas. In males, BMI likely reflects food availability and competitive ability, with individuals that experience food limitation during the breeding season potentially losing mass and displaying microbial shifts consistent with dietary restriction. Another potential factor is anthropogenic feeding by tourists. A large part of the tourism industry in The Bahamas, where high-calorie foods like grapes can elevate blood glucose, disrupt natural foraging patterns, and reduce dietary diversity (Knapp et al., 2013). Over time, these supplements may alter energy storage, body condition, and stress physiology, potentially predisposing iguanas to obesity-like states and immune dysregulation (Ki et al., 2024).

These results indicate that both natural physiological cycles and anthropogenic influences shape variation in alpha diversity. In females that yolk, more follicles show elevated levels of energy metabolites and reactive oxygen species (Webb et al., 2019). Some of our data were collected during the mating season, when these effects are most pronounced and may contribute to observed shifts in the gut microbiome. However, these effects are difficult to disentangle from broader seasonal changes in physiology and available food. Overall, our results suggest that metabolism and general physiology have a significant impact on both individual- and community-level gut microbial diversity unlike reproductive hormones.

## Conclusion

In summary, our study demonstrates that the rock iguana gut microbiome is shaped by a combination of seasonal, reproductive, and physiological factors. Season and sex drive significant shifts in microbial composition, with females showing lower alpha diversity during the breeding season, and a generally unique signature in comparison to males and post-reproductive females, likely due to reproductive demands and associated behavioral and dietary changes. Despite these fluctuations, the core microbiome remains generally stable across seasons and resembles patterns seen in other reptiles, dominated by phyla and families that facilitate plant matter digestion. Energy metabolites, stress markers, and body mass index further influence both alpha- and beta-diversity, in a complex interaction network between physiology, season, and anthropogenic disturbances.

These results confirm previous findings in other lizards, and add to our knowledge about rock iguanas and how external and internal factors affect its microbiome. Future work should explore the mechanisms linking physiological changes with the microbiome, examine seasonal and sex-specific changes longitudinally across multiple breeding cycles, and investigate how rock iguanas retain their core microbiome. More focus should be given to connecting how the energy gets allocated towards reproduction, what role hormones play during the process, and how this reduces the diversity levels in females in particular. Similar focus should be given to confirm if rock iguanas do share the vertical transmission of taxa similar to other species discussed in this study.

## Notes

### Competing Interest Statement

The authors have declared no competing interest.

